# Higher-order intrinsic routes support flexible task-evoked communication

**DOI:** 10.64898/2026.01.28.702230

**Authors:** Garima Bargujar, Siddhartha Singh, Shilpa Dang

## Abstract

Flexible task-evoked communication requires the brain to transiently establish functional interactions that are not evident during rest, yet the network mechanisms supporting these newly formed connections remain unresolved. Using whole-brain functional MRI from 92 participants performing six cognitive tasks, we show that task-evoked communication is implemented within a constrained intrinsic network architecture rather than through large-scale rewiring. Across tasks, we found that approximately 70% of functional connections were preserved from rest, forming a dominant stable core, while task engagement selectively reconfigured a smaller subset of connections, primarily at the between-network level. While existing activity-flow models explain task-evoked information transfer primarily through direct stable core connections, we demonstrate that newly formed connections between regions lacking direct resting-state coupling were supported by selectively strengthened indirect higher-order resting-state routes embedded within the intrinsic scaffold. Network-level analyses revealed a multiscale organization of information transfer: we observed that direct stable routes dominated within-network communication, most prominently in sensory–motor and default mode systems, whereas indirect higher-order routes preferentially supported between-network integration through control and attentional systems. By extending activity-flow modelling to incorporate these higher-order intrinsic routes, we show a significant improvement in the prediction of task-evoked activity at the global, between-network scale, confirming their functional recruitment during task performance. Together, our findings identify higher-order intrinsic routes as a key mechanism enabling flexible task-evoked communication within a stable large-scale network architecture.

## Introduction

The human brain is organized into large-scale functional networks comprising distributed cortical and subcortical regions that exhibit coherent spontaneous activity^1–4^. Interactions among these networks support integrative cognitive processes^3,5,6^. Resting-state functional connectivity (restFC), defined by correlations between regional time series, captures the intrinsic whole-brain organization^7,8^. This topology is constrained by the brain’s structural connectivity scaffold, reflecting synchronous activity patterns shaped, but not fully determined, by white-matter pathways^9–12^. Critically, regions lacking direct anatomical links can still exhibit strong coupling via indirect, higher-order routes mediated through intermediate nodes^13–15^ (**Fig. 1A–B**).

**Figure 1.**
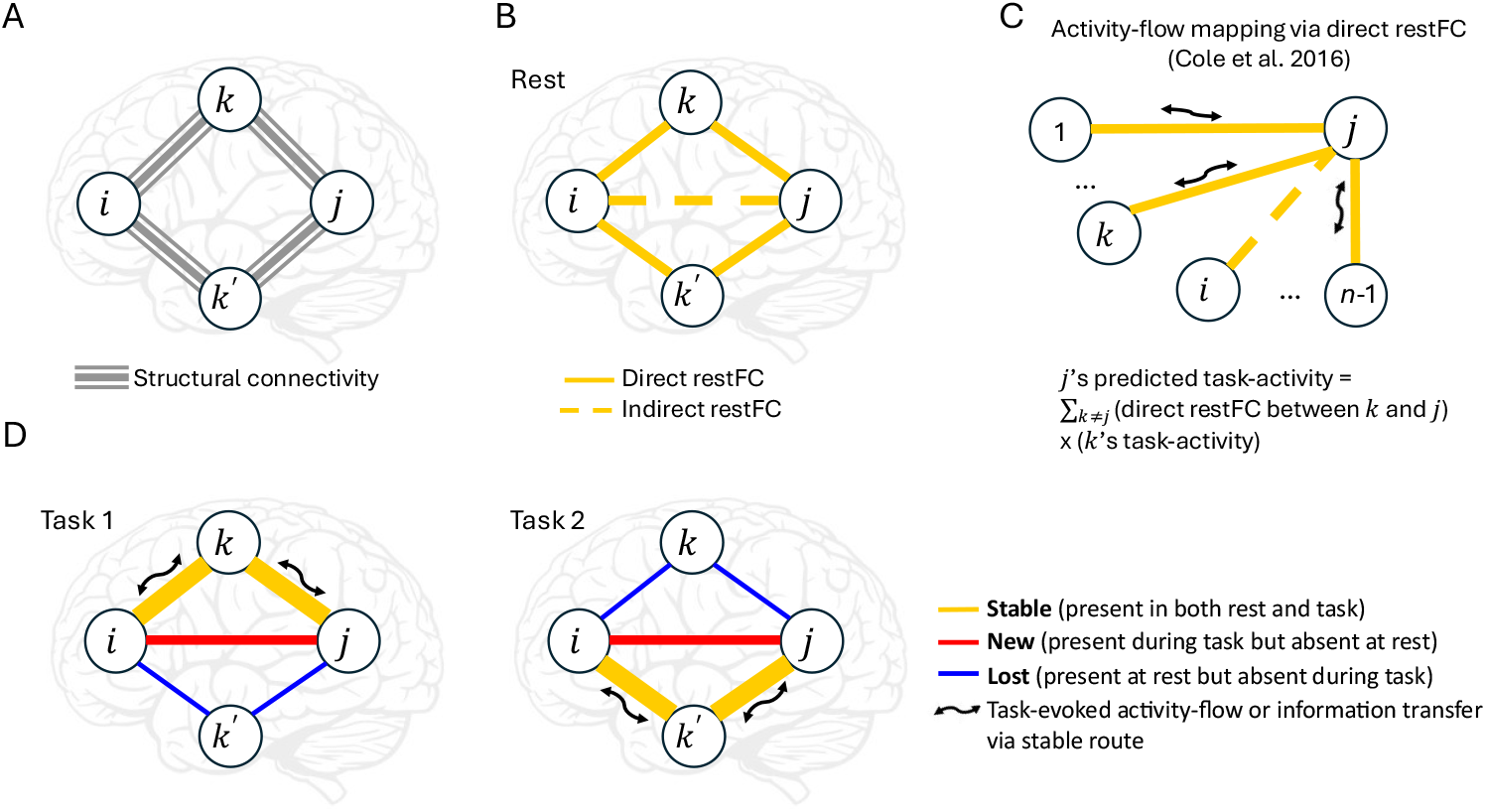
Conceptual framework linking structural connectivity, resting-state functional connectivity (restFC), and task-evoked higher-order information transfer. **(A)** Simplified schematic of structural connectivity between four brain regions, illustrating direct anatomical connections (solid lines) and the absence of direct links between some region pairs. **(B)** Resting-state functional connectivity, derived from spontaneous activity correlations, exhibiting both direct coupling between anatomically connected regions and indirect coupling between regions without direct structural links, mediated via intermediate anatomically connected regions. **(C)** Activity-flow mapping framework^16^, in which task-evoked activity in a target region is predicted from activity in source regions weighted by direct restFC, modelling information transfer along preserved direct resting-state functional routes alone. **(D)** Task engagement induces partial reconfiguration of intrinsic functional connectivity, preserving a stable core (yellow solid lines) while introducing newly formed connections (red solid lines) and suppressing others (blue solid lines). Newly formed task-evoked connections lacking direct restFC route are hypothesized to be supported by selectively strengthened higher-order resting-state routes. Accordingly, distinct task demands are expected to recruit distinct subsets of higher-order routes, enabling flexible yet constrained reconfiguration of information flow.

Accumulating evidence suggests that resting-state network topology serves as an *invariant global routing architecture* supporting task-evoked communication^16,17^. Activity-flow mapping has shown that these intrinsic restFC pathways mechanistically support task-evoked information transfer^16,17^(**Fig. 1C**), and variability in restFC reliably predicts individual and group differences in cognitive performance^18,19^. Complementary evidence from network reconfiguration studies indicate that most task-evoked network topology is preserved from rest (∼80%), with a smaller but systematic task-specific component^20–22^. Task engagement thus reflects a stable intrinsic core accompanied by moderate reconfiguration, expressed through the emergence of novel connections and the selective reweighting (strengthening or suppression) of existing resting-state connections^22^(**Fig. 1D**). These adjustments enable flexible resource allocation and scale with cognitive demand^22^.

Most activity-flow studies have relied on static, linear FC estimates—such as Pearson correlation or multiple regression—implicitly assuming temporally invariant resting-state connectivity. While Pearson correlation is widely used^23,24^, it cannot separate direct dependencies from the indirect ones, leading to false positives. Multiple regression-based FC improves mechanistic specificity by isolating unique regional dependencies and accounting for both common-cause and chain confounds^16,17,25^. Recent extensions have further incorporated causal structure^26^ and probabilistic non-linear routing^27^ to improve prediction robustness. Despite these advances, existing activity-flow frameworks predominantly model task-evoked communication along direct resting-state connections that remain preserved during task performance.

In summary, task engagement entails dynamic reconfiguration layered upon a stable intrinsic core. Yet, existing activity-flow frameworks largely model task-evoked information transfer only along direct resting-state connections that remain preserved during task performance. Consequently, information flow along newly task-evoked connections are not explicitly captured, leaving a critical gap in current models. This raises a fundamental question: how does the brain flexibly implement task-evoked communication while operating within a constrained intrinsic architecture? We propose that novel task-evoked connections between regions lacking direct resting-state coupling do not arise de novo but instead reflect the transient activation (strengthening) of pre-existing indirect, higher-order routes of the intrinsic network scaffold (**Fig. 1D**). Moreover, among the multiple higher-order resting-state routes available between any two regions, task engagement selectively reweights these pathways—strengthening those relevant for current cognitive demands while suppressing alternatives to minimize interference. Under this account, distinct task demands are expected to recruit distinct subsets of higher-order routes, enabling flexible yet constrained reconfiguration of information flow (**Fig. 1D**).

Overall, the proposed framework positions task-evoked network reconfiguration not as a departure from resting-state, but as a selective reweighting of intrinsic global routing architecture, combining core stability with dynamic flexibility. To address the above hypotheses, in this study we sought to systematically characterize how joint modelling of stable and reconfigured connections would support task-evoked information flow. Using publicly available functional magnetic resonance imaging (fMRI) data from 92 healthy adults during resting-state and six cognitive task-states^28^, we first aimed to quantify how task engagement would reorganize whole-brain functional connectivity at rest. We did this by identifying the proportion of connections that were preserved (stable) versus those that were reconfigured (lost and new) during task and determining how these connection types were distributed across local (within-network) and global (between-network) topologies. Building on the central hypothesis that newly formed task connections rely on indirect higher-order stable routes rather than the direct ones; we next examined the multi-step stable core scaffold that would transiently support novel information transfer. Further, to test the functional relevance of this scaffold, we extended the activity-flow modelling framework to incorporate the higher-order stable routes. We evaluated whether these higher-order routes improved prediction of task-evoked activations and whether their contributions differed systematically between local and global communication. Finally, based on above framework, we assessed selective strengthening or weakening of all possible higher-order resting-state routes during task performance and identified task-specific differences. Together, these aims provided a complete framework for understanding how the brain’s stable core scaffold enables flexible, task-specific information transfer via indirect higher-order routes.

Elucidating how intrinsic global routing architecture selectively reweights to support task-evoked information transfer is central to the two overarching goals of the field—one, to establish a unified mechanistic account linking resting-state architecture, task-driven reconfiguration, and activity-flow modelling. Further, this explains how cognitive flexibility is supported within a constrained architecture. Two, to understand how failures of this reweighting process contribute to impaired cognition in brain disorders^29–31^.

## Results

### Mapping Task-Evoked Network Reconfiguration

To quantify how intrinsic brain architecture reorganizes during cognitive tasks, we examined task-evoked network reconfiguration across six tasks—working memory, gambling, relational processing, language, motor, and emotion—using resting-state network topology as the baseline (see *Methods, Pre-processed fMRI Data*). Group-level whole-brain region-to-region binary functional connectivity (FC) was estimated for each task using a multiple linear regression framework applied to mean regional fMRI time series, capturing direct inter-regional dependencies while controlling for shared variance across all other regions and task-evoked co-activation due to externally-induced perturbations^32,33^(see *Methods, Functional Connectivity Estimation*). Importantly, task connectivity was estimated across all brain regions rather than being restricted to task-activated nodes, allowing network reconfiguration to be assessed as task-evoked modulation of large-scale network organization rather than recruitment of isolated regions^34,35^. For visualization, connectivity was summarized at the network level across 11 canonical cortical and subcortical systems (**Fig. 2A–B**). **Panel A** shows network-to-network mean functional connectivity during resting-state, whereas **Panel B** depicts the average across tasks, highlighting preserved core connections alongside newly formed and lost connections during task performance. (For region and network definitions, refer *Methods, Whole-brain ROIs & Networks*.)

**Figure 2.**
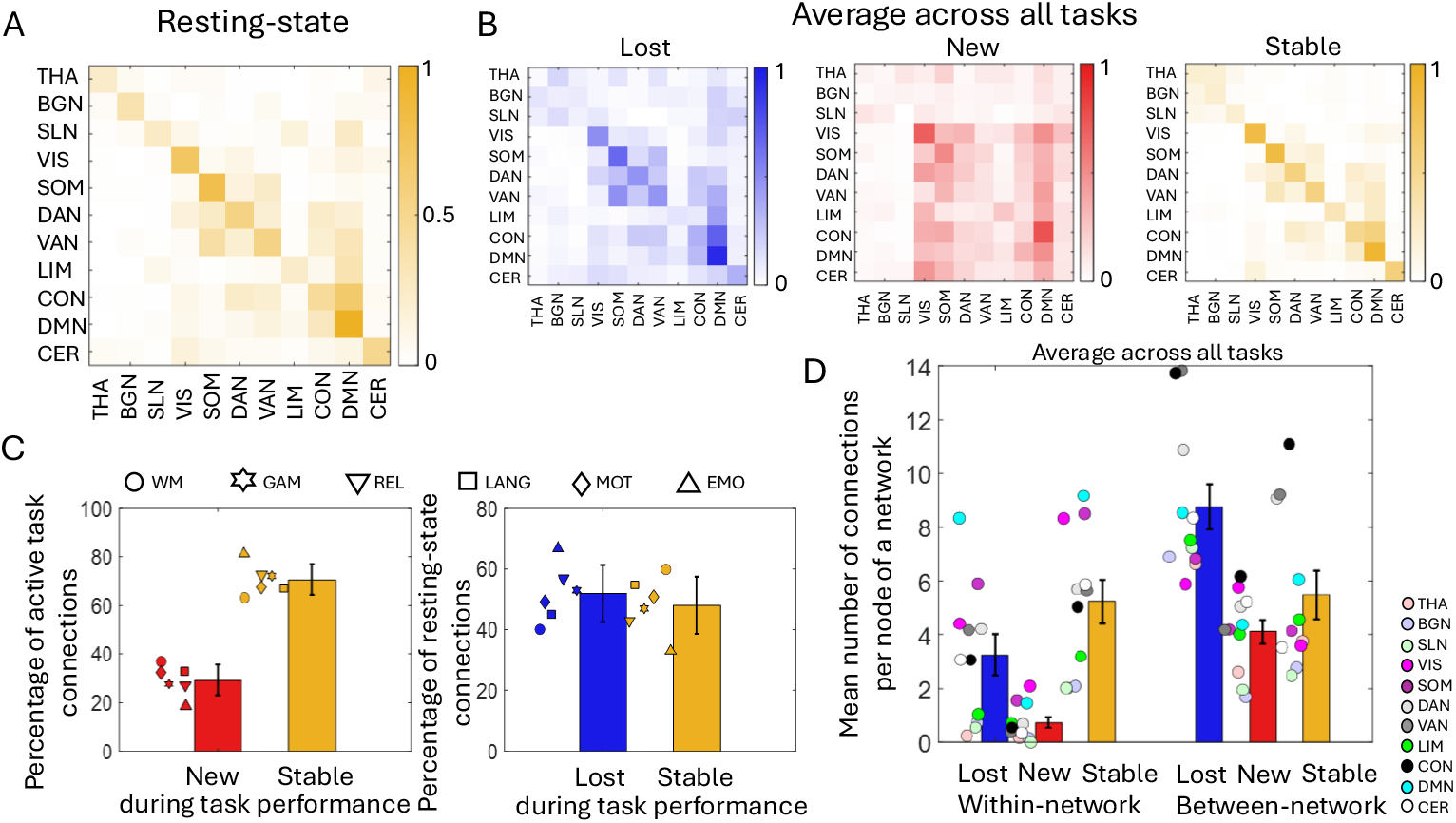
Task-evoked reconfiguration of functional connectivity across large-scale brain networks. **(A)** Group-level network-to-network mean functional connectivity during resting-state, organized according to 11 canonical large-scale networks ^1–3^: thalamic (THA), basal ganglia (BGN), subcortical limbic (SLN), visual (VIS), somatomotor (SOM), dorsal attention (DAN), ventral attention (VAN), limbic (LIM), control (CON), default mode (DMN), and cerebellar (CER) networks. **(B)** Group-level network-to-network mean functional connectivity averaged across all six tasks, shown separately for lost, newly formed, and stable connections and organized by the same network ordering. **(C)** Bar plots showing the percentage of stable and newly formed connections during task performance, as well as lost and stable connections from resting-state, for each task: working memory (WM), gambling (GAM), relational processing (REL), language (LANG), motor (MOT), and emotion (EMO). Bars and error bars represent mean and standard deviation across six tasks. **(D)** Topological distribution of stable and reconfigured connections. The mean number of connections per node shown separately for each connection type (lost, newly formed, and stable) and for within-network and between-network locations. Values represent averages across tasks; task-specific distributions are shown in **Supplementary Fig. S1**. Error bars indicate standard error of mean across 11 networks.

Comparing task and resting-state binary FC revealed a partially reconfigured architecture. Across tasks, 29.32% (±6.33%; mean ± SD) of active task connections were newly formed, whereas the majority (70.68% ± 6.33%) reflected direct stable connections preserved from rest (**Fig. 2C**). Conversely, 51.89% (±9.44%) of resting-state connections were suppressed (lost) during task performance, indicating selective disengagement of task-irrelevant pathways (**Fig. 2C**). These results confirm that task engagement operates on a largely stable intrinsic scaffold while inducing systematic, task-specific reconfiguration^20–22^. Together, these findings establish the coexistence of intrinsic stability and adaptive reconfiguration as a fundamental organizational principle of task-state brain networks.

### Topology of Stability vs. Reconfiguration

Task engagement has been shown to reorganize brain networks primarily by increasing integration across otherwise segregated resting-state communities^36,37^. Building on this view, we examined how task-evoked changes were distributed across two fundamental topological dimensions: within-network (local segregation) and between-network (global integration). For each task, we quantified network-wise mean number of lost, new, and stable connections per node and assessed their distribution across topological locations using repeated-measures ANOVA (see *Methods, ANOVA*).

Across all tasks, we observed the main effects of connection type and location (all *p* < 0.01; **Fig. 2D, Supplementary Fig. S1, and Table S1**), indicating systematic differences in how stability and reconfiguration were organized in the brain. Averaged across tasks, default mode and sensory–motor systems (DMN, SOM, VIS) showed the strongest preservation of within-network stable connections, consistent with a reliance on locally conserved architecture to maintain their unimodal and heteromodal functional specialization during task engagement^38,39^. In contrast, control and attention systems (CON, DAN, VAN) exhibited the highest involvement in between-network connections, encompassing both stable and reconfigured links, highlighting their central role in global coordination across cognitive tasks^17,40,41^. Specifically, when isolating newly formed connections at the between-network level, the highest recruitment was observed in control, visual, and cerebellar systems, indicating that while control and attention networks dominate overall reconfiguration, the emergence of novel task-evoked links additionally engages perceptual and cerebellar networks (**Fig. 2D; Supplementary Fig. S1**).

Critically, a significant connection type × location interaction was observed for every task (all *p* < 0.01; **Supplementary Table S2**). Post-hoc tests revealed that stable core connections were equivalently distributed across within- and between-network levels (all *p* > 0.43), indicating broad preservation of the intrinsic backbone during task performance. By contrast, both newly formed and lost connections were significantly enriched at the between-network level (all *p* < 0.01), demonstrating that task-driven reconfiguration primarily reflects global integrative changes rather than local network restructuring. While this overall topological distribution of stable and reconfigured connections was conserved across tasks, different tasks expressed variance in the relative preservation or reconfiguration of intrinsic connectivity across network locations (**Supplementary Fig. S1**).

Together, these results show that task engagement preserves a widespread stable core across topological dimensions while selectively reconfiguring between-network connectivity, establishing the architectural context in which task-evoked information flow unfolds. We next assessed how this stable and reconfigured topology jointly supported task-evoked information transfer.

### Task-Evoked Information Transfer via Higher-Order Resting-State Routes

In the previous section, we found that across six cognitive tasks approximately 30% of task-evoked connections were newly formed, whereas the remaining 70% reflected a stable core architecture. Because existing activity-flow models capture only task-evoked information transfer via direct stable routes ^16,17^, we reasoned that the information transfer via newly formed task connections — those lacking an underlying direct stable route — might instead occur via indirect higher-order (HO) stable routes embedded within the brain’s intrinsic global communication scaffold. To test whether and how the stable core architecture preserved during a task-state supports newly task-evoked communication, we extended the activity-flow modelling framework^16,17^ by incorporating indirect **higher-order stable routes** identified via the shortest-path computation to predict region-wise task activity (see *Methods, Extended Activity-Flow (EAF) Model*).

Figure 3. summarizes the distribution of indirect HO routes across the large-scale network organization. For each task, we computed the mean number of first-, second-, and third-order routes passing through a network as an intermediate relay (e.g., from VIS to DMN via CON) and averaged these values across tasks. **Panels 3A** and **3B** show all possible HO resting-state routes and the subset preserved during task performance (HO stable routes), respectively, via individual networks. Although defined on resting-state FC, these routes were task-specific because they were identified relative to newly formed edges unique to each task. Within the EAF framework, only stable connections preserved during task states were employed to construct HO routes. HO stable routes constituted a small, selectively strengthened subset of all possible HO resting-state routes (mean ± SD across tasks: 14.64% ± 5.12%) and exhibited substantially greater path strength than lost counterparts (*t*[5] = 17.52, *p* = 10^-5^; mean ± SD: 0.1496 ± 0.0208 vs. 0.0006 ± 0.0002; see *Methods, Identification of higher-order stable routes and path strength*). Path strengths were computed using group-level weighted taskFC. Amongst HO stable routes, the majority were task-specific (60.12% ± 5.32%) and the remainder were shared across (≥2) tasks, i.e., task-general routes. **Fig. 3C** illustrates all possible HO restFC routes and task-general and task-specific HO stable routes expanded node-wise, say from VIS to DMN via CON. Thus, task engagement selectively amplifies a subset of multi-step intrinsic routes, forming an efficient set of higher-order channels that transiently activate newly formed task-evoked connections. This mechanism explains how cognitive flexibility across diverse task demands emerges from the selective recruitment of intrinsic network pathways within a constrained global architecture ^16,20,42,43^.

### Topology of Direct vs. Indirect Stable Routes

To dissect the topology of multi-step communication routes, we next classified each indirect HO stable route as within-network or between-network based on the network membership of the intermediate node. Between-network routes linked nodes in different networks via an intermediate node in a third network (e.g., VIS → DMN via CON), whereas within-network routes involved intermediates belonging to the same network as the endpoints. Further, for each task, we compared network-wise mean number of direct and indirect stable routes per node and assessed their distribution across topological locations (within- and between-networks) using repeated-measures ANOVA, as in the previous topology analysis of stability vs. reconfiguration.

**Figure 3.**
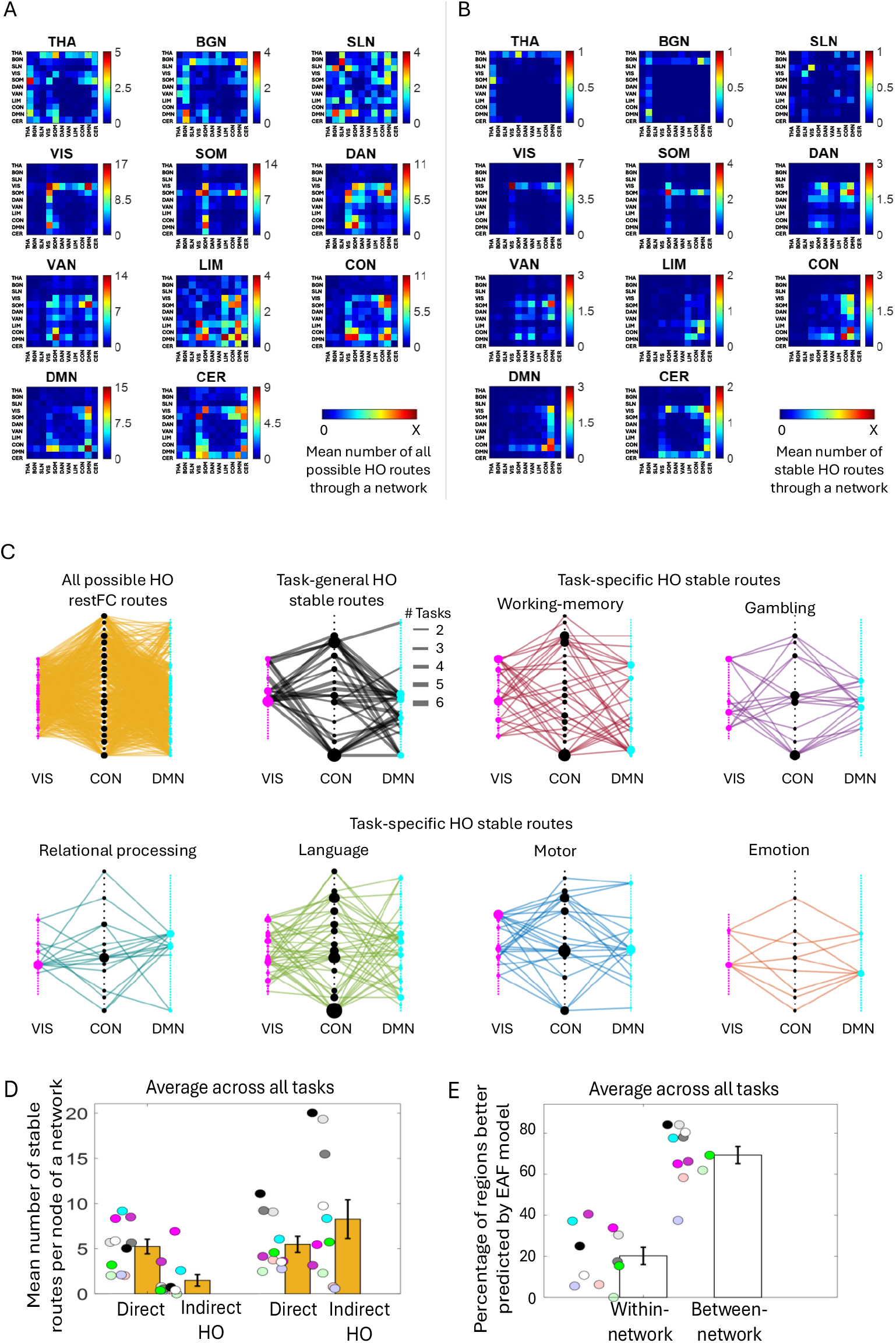
Indirect higher-order resting-state routes supporting flexible task-evoked communication. **(A)** Network-to-network distribution of all possible indirect higher-order (HO) resting-state functional connectivity routes through large-scale brain networks. These multi-step routes link pairs of regions that form newly formed task-evoked connections despite lacking a direct stable restFC edge and are defined by paths traversing one or more intermediate regions within the intrinsic connectivity architecture. **(B)** Network-to-network distribution of the subset of HO routes that are preserved during task performance (HO stable routes). Only a fraction of all possible HO restFC routes is selectively retained and strengthened during task engagement. Values in (A) and (B) are averaged across all tasks. **(C)** Node-wise illustration of all possible HO restFC routes and subset (task-general and task-specific) of HO stable routes, exemplified by connections linking visual (VIS) and default mode (DMN) networks via the control network (CON). Node size is scaled by degree. **(D)** Topological distribution of stable routes, separating direct and indirect HO pathways across two network locations: within-network and between-network. Values represent the mean number of stable routes (direct and indirect) per node, averaged across tasks, and task-specific distributions are shown in **Supplementary Fig. S2. (E)** Contribution of indirect higher-order stable routes to task-evoked information transfer. Shown is the network-wise percentage of regions whose task-evoked activity was better predicted by the extended activity-flow model that incorporates indirect higher-order (HO) routes, evaluated separately at within-network (local) and between-network (global) levels. Values indicate network-wise percentages averaged across tasks; task-specific data is provided in **Supplementary Fig. S3**. In (D) and (E), error bars indicate standard error of mean across 11 networks.

Across all tasks, we observed no main effects of route type or network location (all *p* > 0.01) but a significant route type × location interaction effect, reflecting a systematic crossover in route deployment across topological scales (all *p* < 0.01, **Fig. 3D, Supplementary Fig. S2**, and **Tables S3-S5**). Post hoc comparisons revealed that within networks, direct routes were consistently more prevalent than indirect higher-order routes (mean ± SE: 5.24 ± 0.80 vs. 1.49 ± 0.64, all *p* < 0.01 see **Supplementary Table S4**) and were largely confined to sensory–motor and default mode systems, contributing modestly to local task-evoked processing. In contrast, between-network communication showed the opposite pattern, where indirect HO routes exceeded direct routes (8.26 ± 2.15 vs. 5.48 ± 0.89; although this did not reach significance in all cases, see **Supplementary Table S4**) and were preferentially routed through control and attentional systems. Further, post hoc comparisons revealed that direct route counts were equivalent across within- and between-network locations (5.24 ± 0.80 vs.5.48 ± 0.89, all *p* > 0.44, **Supplementary Table S5**), whereas indirect higher-order routes were significantly more abundant at the between-network level (8.26 ± 2.15 vs. 1.49 ± 0.64, all *p* < 0.02 except for emotion task, **Supplementary Table S5**). While the topological distribution pattern of direct and indirect routes across within- and between-network locations was conserved across tasks, distinct tasks expressed differences in their overall abundance of both direct and higher-order pathways (**Supplementary Fig. S2**). Together, these results indicate a balanced allocation of distinct routing strategies, whereby direct stable routes predominantly support local processing, while indirect higher-order stable routes facilitate global integration during task engagement.

### Validation using the Extended Activity-Flow model

To test whether and how indirect HO routes actively contributed to task-evoked communication, we compared predictions of regional task activity generated by a standard activity-flow model using direct stable routes alone with those generated by the EAF model incorporating both direct and indirect routes. Analyses were performed separately at within- and between-network levels (see *Methods*). This was done to reveal the contribution of indirect higher-order stable routes in information transfer mechanism across both the topological dimensions.

We found that higher-order routes contributed modestly to local information transfer but dominated at the global scale (**Fig. 3E**; **Supplementary Fig. S3**). Within networks, only a subset of regions benefited from inclusion of HO routes (mean ± SE across tasks: 20.22% ± 4.21%), primarily in DMN, VIS, and SOM. This means that at within-network level, task-evoked activity in majority of the regions was better predicted by model 1 including direct routes only. In contrast, at the between-network level, indirect HO routes improved predictions for most regions (69.31% ± 4.18%), with the strongest effects in control and attentional systems, default mode, and cerebellar systems. Paired comparisons across networks were performed for each task, which confirmed significantly greater reliance on HO routes at the between-network than at within-network level for all tasks (all *p* < 0.01; **Supplementary Table S6**). While the overall dominance of higher-order routes for between-network information transfer was conserved across tasks, distinct tasks exhibited systematic differences in the overall proportion of regions relying on these routes (**Supplementary Fig. S3**). Together, these results show that task-evoked information transfer is mediated predominantly by direct resting-state routes locally and by indirect higher-order routes for global integration.

## Discussion

Cognitive flexibility is often attributed to dynamic reconfiguration of information flow in response to changing task demands^44,45^, yet its network-level mechanisms remain unresolved. We show that flexible task-evoked communication is supported by a constrained intrinsic architecture rather than large-scale network rewiring. Across six cognitive tasks, task engagement operated on a dominant stable functional core, with ∼70% of task connections preserved from rest and equivalently expressed across within- and between-network locations. In contrast, a smaller but systematic subset of connections underwent task-specific reconfiguration, predominantly at the between-network level, indicating that cognitive flexibility emerges from selective global reweighting layered upon a stable intrinsic scaffold. While consistent with evidence that resting-state topology provides a persistent backbone for task processing^20–22^, our findings extend this work by identifying the mechanistic routes through which task-evoked connections are functionally implemented. We show that newly formed task connections do not constitute de novo communication channels but are supported by selectively strengthened higher-order routes embedded in the resting-state scaffold, resolving a long-standing limitation of activity-flow models restricted to direct connections^16,17^.

Importantly, task-evoked functional connectivity was corrected for co-activation^32,33^ and estimated at the whole-brain level rather than restricted to strongly task-activated regions. This approach captures task performance as a global network state shaped by distributed interactions—including among weakly activated regions—consistent with evidence that task activation and task connectivity reflect partially dissociable neural processes^34,35^.

Our results reveal a multiscale division of labour. Within networks, information transfer relies on direct intrinsic routes, with sensory–motor and default mode systems showing the strongest expression of these stable pathways. In unimodal sensory–motor networks, direct routes support precise sensory encoding and reliable action execution^39^; in the DMN, they support maintenance and integration of internally generated representations^38^. Between-network transfer is dominated by indirect higher-order routes expressed through control and attentional systems, converging with the integrative hub framework of Ito et al. (2017)^17^, demonstrating that large-scale integration is dynamically implemented via selective task-evoked reweighting.

Functionally, this architecture balances stability and flexibility: a preserved intrinsic core supports fast, low-cost local processing, while higher-order routes are recruited for adaptive integration under complex task demands. Mapping potential routes alone does not establish functional relevance. The Extended Activity-Flow model addresses this gap by predicting regional task activity from direct and indirect pathways. We find a scale-dependent distinction: within-network activity is best predicted by direct routes, whereas including higher-order routes improves predictions at the between-network level. This confirms that higher-order routes are actively recruited during task engagement, bridging the gap between static network topology and dynamic cognitive function^36,37^.

Several limitations warrant consideration. Higher-order routes were estimated using shortest-path approximations on group-level functional connectivity, which may not capture all biologically plausible strategies^46^. Functional connectivity, despite controlling for shared variance across regions, remains an indirect proxy for neural communication. Whole-brain estimation mitigates biases from activation-based node selection, but future work integrating time-resolved measures, causal perturbations, or individual-specific structural connectivity may refine these estimates. Clinically, this architecture suggests that cognitive deficits may arise from impaired route selection or reweighting processes rather than loss of connections, highlighting potential targets for network-based interventions^29–31^.

In summary, flexible task-evoked communication is implemented through higher-order routes embedded within a stable intrinsic network, uniting local processing and large-scale integration. Cognitive flexibility emerges from selective modulation of a preserved global routing scaffold, demonstrating how stability and flexibility are reconciled within a single network framework.

## Methods

### Pre-processed fMRI Data

In this study, we used publicly available pre-processed resting-state functional MRI (rsfMRI) and task fMRI datasets from the Human Connectome Project (HCP)^28,47^. Data were obtained from 92 participants (46 males, 46 females; age range: 22–35 years; mean ± SD: 29.49 ± 3.56 years) drawn from the HCP 100 Unrelated Subjects release. All participants provided written informed consent, and study protocols were approved by the Institutional Review Board at Washington University. All procedures were conducted in accordance with relevant ethical guidelines and regulations. Neuroimaging data were acquired using a Siemens 3T Connectom Skyra scanner equipped with a 32-channel head coil (see HCP documentation for detailed acquisition protocols^48^).

The HCP minimal pre-processing pipeline included correction for gradient nonlinearity, head motion, and EPI distortions, followed by spatial normalization to standard space and intensity normalization. Resting-state fMRI data was further denoised using ICA-FIX, which automatically identifies and removes structured noise components related to motion, physiology, and scanner artifacts^47,49^. No additional temporal filtering or global signal regression was applied beyond the HCP preprocessing pipeline. All further analyses were carried out using custom-written routines in MATLAB software (Version 9.11.0.1809720 (R2021b) Update 1), including functions from Brain Connectivity Toolbox^50^.

#### Resting-state fMRI data

were collected across four scan sessions (14:33 minutes each) over two consecutive days using a gradient-echo echo-planar imaging (EPI) sequence (TR = 720 ms, TE = 33.1 ms, flip angle = 52°, 72 slices, 2 mm slice thickness, 2 mm isotropic voxels, multiband factor = 8). Within each day, oblique axial acquisitions alternated between right-to-left (RL) and left-to-right (LR) phase-encoding directions across successive scan sessions. During scanning, participants were instructed to keep their eyes open and fixate on a centrally presented cross on a dark background.

#### Task fMRI data

Upon completion of the resting-state fMRI scans on each day, participants performed a series of task fMRI paradigms targeting activation of distinct cortical and subcortical systems. Task data were collected across two scanning sessions on separate days, with task order counterbalanced across participants. All tasks were acquired using block or event-related designs optimized for robust activation detection and reliability. Each task consisted of **two scanning sessions per participant**, with identical timing per scan. The task fMRI acquisitions employed identical EPI pulse sequence parameters to those used for resting-state fMRI, except for the task timing duration. Below we present a brief description of individual task paradigms, detailed task protocols can be found elsewhere^51^:

#### Working Memory (WM)

Participants performed a block-design N-back working memory task involving visual stimuli drawn from four categories: places, tools, faces, and body parts (non-mutilated, with no nudity)^52^. Within each scanning session, a total of eight stimulus blocks (25 s each) were presented interleaved with four fixation blocks (15 s each) and stimulus categories were shown in separate blocks. Half of the blocks employed a 2-back working memory condition, and half employed a 0-back condition, which served as a control comparison. At the beginning of each block, a 2.5 s instruction cue indicated the task condition and, for 0-back blocks, the target stimulus. A block consisted of 10 trials. During each trial, stimuli were presented for 2 s, followed by a 500 ms inter-trial interval (ITI).

#### Gambling (GAM)

The gambling task was adapted from a previous study^53^ and involved a card-guessing paradigm designed to probe reward and loss processing. Within each scanning session, trials were organized into blocks of eight trials that were either predominantly reward or predominantly loss, with trial types pseudo-randomly interleaved within each block (e.g. in case of reward, 6 reward trials were pseudo-randomly interleaved with either 1 neutral and 1 loss trial, 2 neutral trials, or 2 loss trials). Each session comprised two mostly reward blocks and two mostly loss blocks (28 s each), interleaved with four 15 s fixation blocks. On each trial, participants indicated whether the value of a hidden card (range: 1–9) was higher or lower than five using a button response. The decision cue (“?”) was presented for up to 1.5 s, followed by outcome feedback for 1 s and a 1 s inter-trial interval. Feedback signalled reward (+$1), loss (−$0.50), or neutral outcomes using color-coded visual cues.

#### Relational Processing (REL)

The relational processing task assessed higher-order relational reasoning using visual stimuli comprising shapes filled with textured patterns, adapted from a previous study^54^. Each scanning session contained three relational blocks (18 s each), three matching blocks (18 s each), and three fixation blocks (16 s each). In the relational condition, participants viewed two pairs of objects and first determined the dimension along which the top pair differed (shape or texture), then judged whether the bottom pair differed along the same dimension. In the relational block, consisting of four trials, stimuli were presented for 3.5 s with a 500 ms inter-trial interval. In the control matching condition, participants matched a single bottom object to one of two top objects based on a cued dimension (shape or texture). Responses were binary (yes/no). In the matching block, consisting of five trials, stimuli were presented for 2.8 s with a 400 ms inter-trial interval.

#### Language (LANG)

In the language task, developed by Binder et al. ^55^, blocks of story comprehension alternated with blocks of arithmetic processing. Each scanning session included four story blocks and four math blocks, with block durations varying but averaging approximately 30 s; math blocks were matched in length to the story blocks, with additional math trials appended as needed to complete the ∼3.8-min session. During story blocks, participants listened to brief auditory narratives (5–9 sentences) adapted from Aesop’s fables, followed by a two-alternative forced-choice question probing story content. During math blocks, participants solved aurally presented addition and subtraction problems, selecting the correct answer from two alternatives via button press. Task difficulty in the math condition was adaptively adjusted to maintain comparable performance across participants.

#### Motor (MOT)

The motor task, adapted from a previous study ^2^, was designed to map somatotopic motor representations. Each scanning session included ten 15 s movement blocks (two tongue blocks, four hand blocks (two left, two right), and four foot blocks (two left, two right)) as well as three 15 s fixation blocks. Participants followed visual cues instructing them to perform movements of the left or right fingers, left or right toes, or the tongue. Each movement lasted 12 s, preceded by a 3-s instruction cue.

#### Emotion (EMO)

The emotion task, adapted from previous works by Hariri et al. ^56,57^, assessed affective processing using a face-matching paradigm. Each scanning session comprised three face blocks (21 s each) and three shape blocks (21 s each), with an additional 8 s of fixation at the end of each session. Participants were instructed to select which of two stimuli presented at the bottom of the screen matched a target stimulus displayed at the top. In face blocks, the stimuli were faces expressing angry or fearful emotions, whereas in shape blocks participants matched geometric shapes. Trials were organized into blocks of six trials of the same condition (face or shape). Each stimulus was presented for 2 s, followed by a 1 s inter-trial interval. Blocks were preceded by a 3 s instruction cue (“face” or “shape”), yielding a total block duration of 21 s.

### Whole-brain Regions-of-interest & Networks

We adopted a multimodal parcellation strategy to define whole-brain regions of interest (ROIs). Cortical parcellation comprised 400 regions (200 per hemisphere) derived from a homotopic atlas^58^, complemented by 26 cerebellar regions from the Automated Anatomical Labelling 3 (AAL3) atlas^59^. Subcortical structures were delineated into 54 parcels (27 per hemisphere) using a multiscale subcortical atlas^60^. Together, these schemes yielded a total of 480 ROIs.

The 426 cortical and cerebellar parcels were organized into eight canonical resting-state networks: visual (VIS), somatomotor (SOM), dorsal attention (DAN), ventral attention (VAN), limbic (LIM), control (CON), default mode (DMN), and cerebellar (CER) networks^1,2^. In parallel, the 54 subcortical parcels were assigned to three recently characterized subcortical resting-state networks: thalamic (THA), basal ganglia (BGN), and subcortical limbic (SLN) networks^3^.

### Functional Connectivity Estimation

For each participant, mean regional time series were extracted from pre-processed resting-state and task fMRI data for all regions (*N* = 480, refer to previous section). To minimize inter-session baseline differences, regional time series from individual scanning sessions were mean-centred prior to concatenation across sessions to generate a “representative” time series for each region.

### Whole-brain Resting-state Functional Connectivity

Resting-state functional connectivity was estimated using a whole-brain multiple linear regression framework^61^. For each target region, its representative time series was modelled as a linear combination of the time series from all other regions, which served as predictors or regressors. Regression coefficients were estimated using ordinary least squares, yielding a directed set of weights that quantify the unique contribution of each source region to variance in the target region’s activity after accounting for all remaining regions. For each target region *j*, the regression model was defined as:

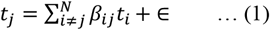

where, *t*_*i*_ and *t*_*j*_ denote the representative time series for regions *i* and *j*, respectively, *β*_*ij*_ represents the directed FC weight from source region *i* to target region *j*, ∈ denotes the residual error, and *i* = 1 *to N* but *i* ≠ *j*, such that the number of predictor regions is *N-1* (= 479). This procedure yielded region-to-region FC estimates that captured the unique explanatory influence of each source region on the target region’s activity, rather than shared variance across regions. Because all regional time series were mean-centred, the regression models were fit without an intercept term. This regression-based FC estimation was performed independently for each participant and repeated for every region across the whole brain.

### Whole-brain Task-evoked Functional Connectivity

Whole-brain functional connectivity during task performance was estimated using a multivariate linear regression framework applied to pre-processed task fMRI time series. For each target region, its representative time series was modelled as a linear function of the time series from all other regions, like equation (1) above. In addition, we explicitly included a convolved task regressor corresponding to the experimental design. The task regressor was defined as a binary vector taking a value of 1 during task blocks and 0 during fixation (baseline) blocks and further convolved with a canonical hemodynamic response function. Including task regressors is essential to prevent task-locked co-activation among regions from being misattributed to inter-regional coupling. This procedure minimizes inflation of connectivity estimates due to task-evoked co-activation^32,33^. By conditioning simultaneously on activity in all other regions and task design modulation, this approach yielded unique, co-activation controlled, region-to-region connectivity estimates during task performance.

Furthermore, classic task fMRI analyses emphasize task-evoked activation or connectivity restricted to task-activated regions, effectively focusing on task activity relative to rest^62,63^. While informative for identifying engaged regions, this approach conflates activation with interaction and overlooks network-wide changes beyond strongly activated nodes^34,35^. In contrast, we treated task performance as a *global network state*, enabling direct comparison between whole-brain intrinsic connectivity present at rest and reconfiguration during task execution. Estimating task-evoked functional connectivity at the whole-brain level allows us to quantify how much inter-regional coupling is preserved from resting-state architecture versus selectively reconfigured under task demands, framing task performance as modulation of large-scale network organization rather than recruitment of isolated regions.

### Group-level Whole-brain Functional Connectivity Matrices

*Group-level inference* was performed, separately for both rest and task fMRI data, by subjecting the respective regression coefficients obtained for each participant to one-sample *t*-tests across participants, constituting a random-effects analysis^64^. Statistical significance was assessed using a threshold of *p* < 0.05, Bonferroni-corrected for multiple comparisons, with the correction factor equal to the number of predictors in the model to account for the partial dependence among regional signals^65,66^.

Based on these group-level statistics, we constructed three whole-brain functional connectivity matrices: Firstly, a region-to-region binary connectivity matrix ***B*** = [*b*_*ij*_] was defined, in which an entry *b*_*ij*_ was set to 1 if activity in a source region *i* exerted statistically significant influence on activity in a target region *j*, and 0 otherwise. Secondly, a region-to-region weighted connectivity matrix ***W*** = [*β*_*ij*_] simply reflected the group-level regression coefficients between the regions. Lastly, we also computed a group-level whole-brain network-to-network mean connectivity matrix using the region-to-region binary connectivity ***B***. For this, we computed the mean number of connections 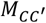from the source nodes *i* ∈ *C* to target nodes *j* ∈ *C*′ as follows:

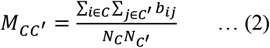

where, *b*_*ij*_ is the group-level binary FC from node *i* to *j, N*_*c*_ and 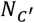 are the number of nodes in network *C* and *C*′, respectively. For task fMRI data, the whole-brain connectivity matrices were computed separately for the three distinct connection types, i.e. lost, new and stable.

### ANOVA

To quantify how task engagement modulates the topological distribution of functional connectivity across the brain’s intrinsic network architecture, we performed a repeated-measures analysis of variance (ANOVA) separately for each task. Task-related configuration changes were defined by comparing group-level whole-brain binary FC matrices derived from task and resting-state data. Task connections were classified as stable (present in both rest and task), new (present only during task), or lost (present only during rest). Each connection was further categorized as within-network or between-network based on membership according to 11 canonical resting-state networks^1–3^.

For each network *C*, we computed the mean number of connections per node separately for each combination of connection type (lost, new, stable) and topological location (within-vs. between-network). Specifically, mean within-network 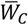 and between-network 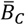 connectivity measures were quantified by summing binary connections from each source node *i* ∈ *C* to target nodes inside (*j* ∈ *C*) or outside (*j* ∉ *C*) the same network, respectively, and averaging across all nodes in that network:

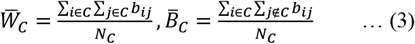

where, *b*_*ij*_ denotes the group-level binary FC from node *i* to node *j*, and *N*_*C*_ is the number of nodes in network *C*.

These mean connectivity measures were entered into a two-way repeated-measures ANOVA with connection type (3 levels: stable, new, lost) and location (2 levels: within-network, between-network) as within-design factors, and network identity (11 levels) as the repeated measure. This analysis assessed the main effects of connection type and location, as well as their interaction, to determine whether task-evoked reconfiguration preferentially targeted local versus global network organization. When significant effects were detected, Bonferroni-corrected post hoc tests were performed to assess pairwise differences between factor levels. Exact F-statistics and corresponding p-values are reported in supplementary information.

### Identification of Higher-order Stable Routes and Path Strength

For each task, we first identified the newly formed task-evoked functional connections—edges that were not directly present in the group-level binary restFC matrix. Then, using the group-level binary restFC we computed the shortest paths between all pairs of regions exhibiting newly formed connections during a task-state. This yielded *the set of all possible higher-order (multi-step) restFC routes* that connected these region pairs through intermediate nodes, forming the basis of potential task-evoked information transfer via indirect restFC pathways. We computed such higher-order paths up till third order (3 intermediate nodes) because the characteristic path length^67^ across resting-state connectivity matrix was found to be 2.19 (±0.64). For a region pair with newly formed taskFC, to start first-order restFC routes were identified as no direct stable connection (restFC) was present between them and subsequently, higher-order routes were identified in case no lower-order route was found between the pair.

Thereafter, we tested whether all possible HO resting-state routes existed during a task-state or not. Those that existed were termed as the HO stable routes. Further, we also tested whether there was a significant difference between the path strengths of stable vs lost HO routes during the tasks. For this, we compared their mean path strengths for the six tasks using a paired t-test. The path strength was computed using the weighted taskFC matrix, i.e. by multiplying the individual weights of connections forming that path.

### Extended Activity-Flow Model

To evaluate the extent to which the stable core connections preserved during a task-state supported task-evoked information transfer, we developed an *extended activity-flow (EAF) modelling* framework to include the *higher-order stable routes*. Following Cole et al. (2016), regional task-evoked activity was predicted as the weighted sum of activity in connected regions. Predicted time series for individual regions were obtained separately for each task and participant. The EAF model was estimated separately at local and global levels to reveal the distribution of higher-order stable routes across two fundamental topological dimensions. Two nested models were implemented separately at both *within-network (local)* and *between-network (global)* levels as follows:

#### Model 1 (Direct-only model)

Predictions were derived solely using the connections at rest which existed during task, i.e. the direct stable routes^16,17^, as shown below:

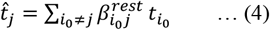

Where 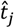and *t*_*j*_ are the predicted and actual task-activities of region *j*, respectively and 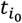 is the actual task-activity of region *i*_0_. In this model, we consider only those source regions *i*_0_ which have direct stable routes along with their resting-state FC weight to target region *j*, denoted by 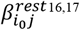.

#### Model 2 (Direct + Indirect HO model)

Predictions were derived using both the direct stable routes and the newly identified indirect higher-order stable routes for a target region, as shown below:

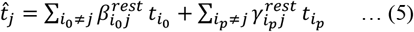

Where 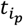 is the actual task-activity of source region *i*_*p*_, which has indirect *p*-th order stable routes to target region *j*, where *p* ∈ {1, 2, 3}. Since multiple *p*-th order routes can exist between a pair of regions (e.g., *i*_*p*_ and *j*) via multiple intermediate nodes, we defined the total strength between the regions as the sum of individual path-strengths as 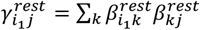 for first-order routes, as 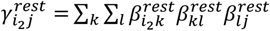 for second-order routes, and so on for third-order routes.

For each task, participant, and network-level, model performance was evaluated by computing the residual sum of squares (RSS) between the predicted and actual regional time series. For each region j, we computed *F-statistic* comparing RSS_1_ (Model 1) and RSS_2_ (Model 2):

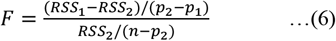

where *p*_1_ and *p*_2_ are the number of parameters in models 1 and 2, respectively, and *n* is the number of time points. A stronger positive F-statistic value indicates that model 2 explains the variance in task-activity of region *j* significantly better than model 1.

### Group-level and network-wise summary statistics

For each task and network-level, we performed a group-level random effects analysis^64^ over region-specific F-statistic values (say, region *j*) from all participants. Significant F-statistic at group-level (*p* < 0.05, Bonferroni corrected for multiple comparisons) identified regions for which inclusion of indirect higher-order stable routes (model 2) led to significantly better prediction accuracy than direct stable routes only (model 1). Further, we computed for each of the 11 canonical brain networks, the percentage of regions better predicted by model 2 than model 1 (significantly positive *F*-statistic across participants). This yielded network-wise measure of local (within-network) versus global (between-network) reliance on higher-order pathways.

## Authors’ contributions

GB – Conceptualization, all main analyses including ANOVA, topology analysis, and EAF modelling, writing first draft, reviewing and editing manuscript. SS – data curation and functional connectivity estimation. SD – Conceptualization, reviewing and editing manuscript.

## Acknowledgements

The authors would like to thank bachelor students Chinmay Vashishth and Neermita Bhattacharya for their help with task fMRI data download and time series extraction.

## Data and code availability

The datasets used in the current study are publicly available from Human Connectome Project. The source data and source code (.m) files used for analysis and generation of results in the current study can be obtained from the corresponding author upon a reasonable request.

## Notes

**<u>Conflict of interests:</u>** The authors declare no competing interests.

https://www.humanconnectome.org/study/hcp-young-adult

### Competing Interest Statement

The authors have declared no competing interest.

## References

1. Thomas Yeo, B. T. et al. The organization of the human cerebral cortex estimated by intrinsic functional connectivity. J. Neurophysiol. 106, 1125–1165 (2011).

2. Buckner, R. L., Krienen, F. M., Castellanos, A., Diaz, J. C. & Thomas Yeo, B. T. The organization of the human cerebellum estimated by intrinsic functional connectivity. J.Neurophysiol. 106, 2322–2345 (2011).

3. Jallepalli, D. & Dang, S. Cortico-subcortical converging organization at rest. Sci. Rep. 15, (2025).

4. Fox Michael D & Raichle Marcus E. Spontaneous fluctuations in brain activity observed with functional magnetic resonance imaging. Nat. Rev. Neurosci. 8, 700–711 (2007).

5. Wang, R. et al. Segregation, integration, and balance of large-scale resting brain networks configure different cognitive abilities. Proc. Natl. Acad. Sci. U. S. A. 118, (2021).

6. Bell, P. T. & Shine, J. M. Estimating Large-Scale Network Convergence in the Human Functional Connectome. Brain Connect. 5, 565–574 (2015).

7. Biswal, B. B. & Uddin, L. Q. The history and future of resting-state functional magnetic resonance imaging. Nature 641, 1121–1131 (2025).

8. Liu, Z. Q. et al. Benchmarking methods for mapping functional connectivity in the brain. Nat. Methods 22, 1593–1602 (2025).

9. Hermundstad, A. M. et al. Structural foundations of resting-state and task-based functional connectivity in the human brain. Proc. Natl. Acad. Sci. U. S. A. 110, 6169–6174 (2013).

10. Honey, C. J. et al. Predicting human resting-state functional connectivity from structural connectivity. Proc. Natl. Acad. Sci. U. S. A. 106, 2035–2040 (2009).

11. Fotiadis, P. et al. Structure–function coupling in macroscale human brain networks. Nat. Rev. Neurosci. (2024) doi:10.1038/s41583-024-00846-6.

12. Collins, E. et al. Mapping the structure-function relationship along macroscale gradients in the human brain. Nat. Commun. 15, (2024).

13. de Pasquale, F., Della Penna, S., Sabatini, U., Caravasso Falletta, C. & Peran, P. The anatomical scaffold underlying the functional centrality of known cortical hubs. Hum. Brain Mapp. 38, 5141–5160 (2017).

14. Skudlarski, P. et al. Measuring brain connectivity: Diffusion tensor imaging validates resting state temporal correlations. Neuroimage 43, 554–561 (2008).

15. Van Den Heuvel, M. P., Mandl, R. C. W., Kahn, R. S. & Hulshoff Pol, H. E. Functionally linked resting-state networks reflect the underlying structural connectivity architecture of the human brain. Hum. Brain Mapp. 30, 3127–3141 (2009).

16. Cole, M. W., Ito, T., Bassett, D. S. & Schultz, D. H. Activity flow over resting-state networks shapes cognitive task activations. Nat. Neurosci. 19, 1718–1726 (2016).

17. Ito, T. et al. Cognitive task information is transferred between brain regions via resting-state network topology. Nat. Commun. 8, (2017).

18. Zhang, C. et al. Resting-state BOLD signal variability is associated with individual differences in metacontrol. Sci. Rep. 12, (2022).

19. Millar, P. R. et al. Evaluating cognitive relationships with resting-state and task-driven blood oxygen level-dependent variability. J. Cogn. Neurosci. 33, 279–302 (2020).

20. Cole, M. W., Bassett, D. S., Power, J. D., Braver, T. S. & Petersen, S. E. Intrinsic and task-evoked network architectures of the human brain. Neuron 83, 238–251 (2014).

21. Krienen, F. M., Thomas Yeo, B. T. & Buckner, R. L. Reconfigurable task-dependent functional coupling modes cluster around a core functional architecture. Philos. Trans. R. Soc. B Biol. Sci. 369, (2014).

22. Zhang, W., Tang, F., Zhou, X. & Li, H. Dynamic Reconfiguration of Functional Topology in Human Brain Networks: From Resting to Task States. Neural Plast. 2020, (2020).

23. Smith, S. M. et al. Network modelling methods for FMRI. Neuroimage 54, 875–891 (2011).

24. Zalesky, A., Fornito, A. & Bullmore, E. On the use of correlation as a measure of network connectivity. Neuroimage 60, 2096–2106 (2012).

25. Reid, A. T. et al. Advancing functional connectivity research from association to causation. Nat. Neurosci. 22, 1751–1760 (2019).

26. Sanchez-Romero, R., Ito, T., Mill, R. D., Hanson, S. J. & Cole, M. W. Causally informed activity flow models provide mechanistic insight into network-generated cognitive activations. Neuroimage 278, (2023).

27. Zhu, H. et al. Activity flow mapping over probabilistic functional connectivity. Hum. Brain Mapp. 44, 341–361 (2023).

28. Smith, S. et al. The WU-Minn Human Connectome Project: An overview. Neuroimage 80, 62–79 (2013).

29. Frisoni, G. B., Pievani, M., De Haan, W., Wu, T. & Seeley, W. W. Functional network disruption in the degenerative dementias. Lancet Neurol. 10, 829–843 (2011).

30. Hearne, L. J. et al. Activity flow underlying abnormalities in brain activations and cognition in schizophrenia. Sci. Adv. 7, (2021).

31. Wang, M. et al. Disrupted dynamic network reconfiguration of the brain functional networks of individuals with autism spectrum disorder. Brain Commun. 4, (2022).

32. Gerchen, M. F., Bernal-Casas, D. & Kirsch, P. Analyzing task-dependent brain network changes by whole-brain psychophysiological interactions: A comparison to conventional analysis. Hum. Brain Mapp. 35, 5071–5082 (2014).

33. Gerchen, M. F. & Kirsch, P. Combining task-related activation and connectivity analysis of fMRI data reveals complex modulation of brain networks. Hum. Brain Mapp. 38, 5726–5739 (2017).

34. Di, X. & Biswal, B. B. Toward Task Connectomics: Examining Whole-Brain Task Modulated Connectivity in Different Task Domains. Cereb. Cortex 29, 1572–1583 (2019).

35. Masharipov, R., Knyazeva, I., Korotkov, A., Cherednichenko, D. & Kireev, M. Comparison of whole-brain task-modulated functional connectivity methods for fMRI task connectomics. Commun. Biol. 7, (2024).

36. Hearne, L. J., Cocchi, L., Zalesky, A. & Mattingley, J. B. Reconfiguration of brain network architectures between resting-state and complexity-dependent cognitive reasoning. J.Neurosci. 37, 8399–8411 (2017).

37. Crossley, N. A. et al. Cognitive relevance of the community structure of the human brain functional coactivation network. Proc. Natl. Acad. Sci. U. S. A. 110, 11583–11588 (2013).

38. Paquola, C. et al. The architecture of the human default mode network explored through cytoarchitecture, wiring and signal flow. Nat. Neurosci. 28, 654–664 (2025).

39. Lee, W. H. & Frangou, S. Linking functional connectivity and dynamic properties of resting-state networks. Sci. Rep. 7, (2017).

40. Cole, M. W., Pathak, S. & Schneider, W. Identifying the brain’s most globally connected regions. Neuroimage 49, 3132–3148 (2010).

41. van den Heuvel, M. P. & Sporns, O. Network hubs in the human brain. Trends Cogn. Sci. 17, 683–696 (2013).

42. Cole, M. W., Etzel, J. A., Zacks, J. M., Schneider, W. & Braver, T. S. Rapid Transfer of Abstract Rules to Novel Contexts in Human Lateral Prefrontal Cortex. Front. Hum. Neurosci. 5, (2011).

43. Shine, J. M. & Poldrack, R. A. Principles of dynamic network reconfiguration across diverse brain states. Neuroimage 180, 396–405 (2018).

44. Braun, U. et al. Dynamic reconfiguration of frontal brain networks during executive cognition in humans. Proc. Natl. Acad. Sci. U. S. A. 112, 11678–11683 (2015).

45. Uddin, L. Q. Cognitive and behavioural flexibility: neural mechanisms and clinical considerations. Nat. Rev. Neurosci. 22, 167–179 (2021).

46. Neudorf, J., Kress, S. & Borowsky, R. Comparing models of information transfer in the structural brain network and their relationship to functional connectivity: diffusion versus shortest path routing. Brain Struct. Funct. 228, 651–662 (2023).

47. Glasser, M. F. et al. The minimal preprocessing pipelines for the Human Connectome Project. Neuroimage 80, 105–124 (2013).

48. Van Essen, D. C. et al. The Human Connectome Project: A data acquisition perspective. Neuroimage 62, 2222–2231 (2012).

49. Griffanti, L. et al. ICA-based artefact removal and accelerated fMRI acquisition for improved resting state network imaging. Neuroimage 95, 232–247 (2014).

50. Rubinov, M., Kötter, R., Hagmann, P. & Sporns, O. Brain connectivity toolbox: a collection of complex network measurements and brain connectivity datasets. Neuroimage 47, S169 (2009).

51. Barch, D. M. et al. Function in the human connectome: Task-fMRI and individual differences in behavior. Neuroimage 80, 169–189 (2013).

52. Drobyshevsky, A., Baumann, S. B. & Schneider, W. A rapid fMRI task battery for mapping of visual, motor, cognitive, and emotional function. Neuroimage 31, 732–744 (2006).

53. Delgado, M. R., Nystrom, L. E., Fissell, C., Noll, D. C. & Fiez, J. A. Tracking the hemodynamic responses to reward and punishment in the striatum. J. Neurophysiol. 84, 3072–3077 (2000).

54. Smith, R., Keramatian, K. & Christoff, K. Localizing the rostrolateral prefrontal cortex at the individual level. Neuroimage 36, 1387–1396 (2007).

55. Binder, J. R. et al. Mapping anterior temporal lobe language areas with fMRI: A multicenter normative study. Neuroimage 54, 1465–1475 (2011).

56. Hariri, A. R. et al. Preference for immediate over delayed rewards is associated with magnitude of ventral striatal activity. J. Neurosci. 26, 13213–13217 (2006).

57. Hariri, A. R., Tessitore, A., Mattay, V. S., Fera, F. & Weinberger, D. R. The amygdala response to emotional stimuli: A comparison of faces and scenes. Neuroimage 17, 317–323 (2002).

58. Yan, X. et al. Homotopic local-global parcellation of the human cerebral cortex from resting-state functional connectivity. Neuroimage 273, (2023).

59. Rolls, E. T., Huang, C. C., Lin, C. P., Feng, J. & Joliot, M. Automated anatomical labelling atlas 3. Neuroimage 206, (2020).

60. Tian, Y., Margulies, D. S., Breakspear, M. & Zalesky, A. Topographic organization of the human subcortex unveiled with functional connectivity gradients. Nat. Neurosci. 23, 1421–1432 (2020).

61. Wang, Y., Kang, J., Kemmer, P. B. & Guo, Y. An efficient and reliable statistical method for estimating functional connectivity in large scale brain networks using partial correlation. Front. Neurosci. 10, (2016).

62. Dolan, R. J. et al. Psychophysiological and modulatory interactions in neuroimaging. Neuroimage 6, 218–229 (1997).

63. Friston, K. J. Functional and Effective Connectivity: A Review. Brain Connect. 1, 13–36 (2011).

64. Penny, W. & Holmes, A. Random-Effects Analysis. in Human Brain Function: Second Edition 843–850 (2003). doi:10.1016/B978-012264841-0/50044-5.

65. Mheich, A., Wendling, F. & Hassan, M. Brain network similarity: Methods and applications. Netw. Neurosci. 4, 507–527 (2020).

66. Rice, W. R. Analyzing Tables of Statistical Tests. Evolution (N. Y). 43, 223 (1989).

67. Sporns, O. Network attributes for segregation and integration in the human brain. Curr. Opin. Neurobiol. 23, 162–171 (2013).

