## Supplementary Information for "Higher-order intrinsic routes support flexible task-evoked communication"

#### Task-specific topology of stable and reconfigured connections

Extending the task-related reconfiguration patterns described in main results (**Fig. 2**), we examined whether individual tasks differed in how they redistributed stable, newly formed, and lost connections across within- and between-network levels (**Fig. S1**). Working memory task exhibited the highest mean number of stable connections together with newly formed connections, and the lowest degree of connection loss relative to rest. This pattern was observed at both within-network and between-network scales, indicating substantial preservation of intrinsic connectivity alongside selective task-related augmentation.

In contrast, the emotion task showed a distinct reconfiguration profile, characterized by the lowest mean number of stable and newly formed connections and the highest mean number of lost connections across network levels, reflecting stronger suppression of resting-state connectivity. Other tasks, including gambling, relational processing, language, and motor tasks exhibited intermediate profiles, with moderate preservation of stable connections, partial recruitment of new connections, and moderate connection loss. Together, these results indicate that while the relative balance of stability and reconfiguration was conserved across tasks, tasks differed systematically in their strengths of network preservation and reconfiguration.

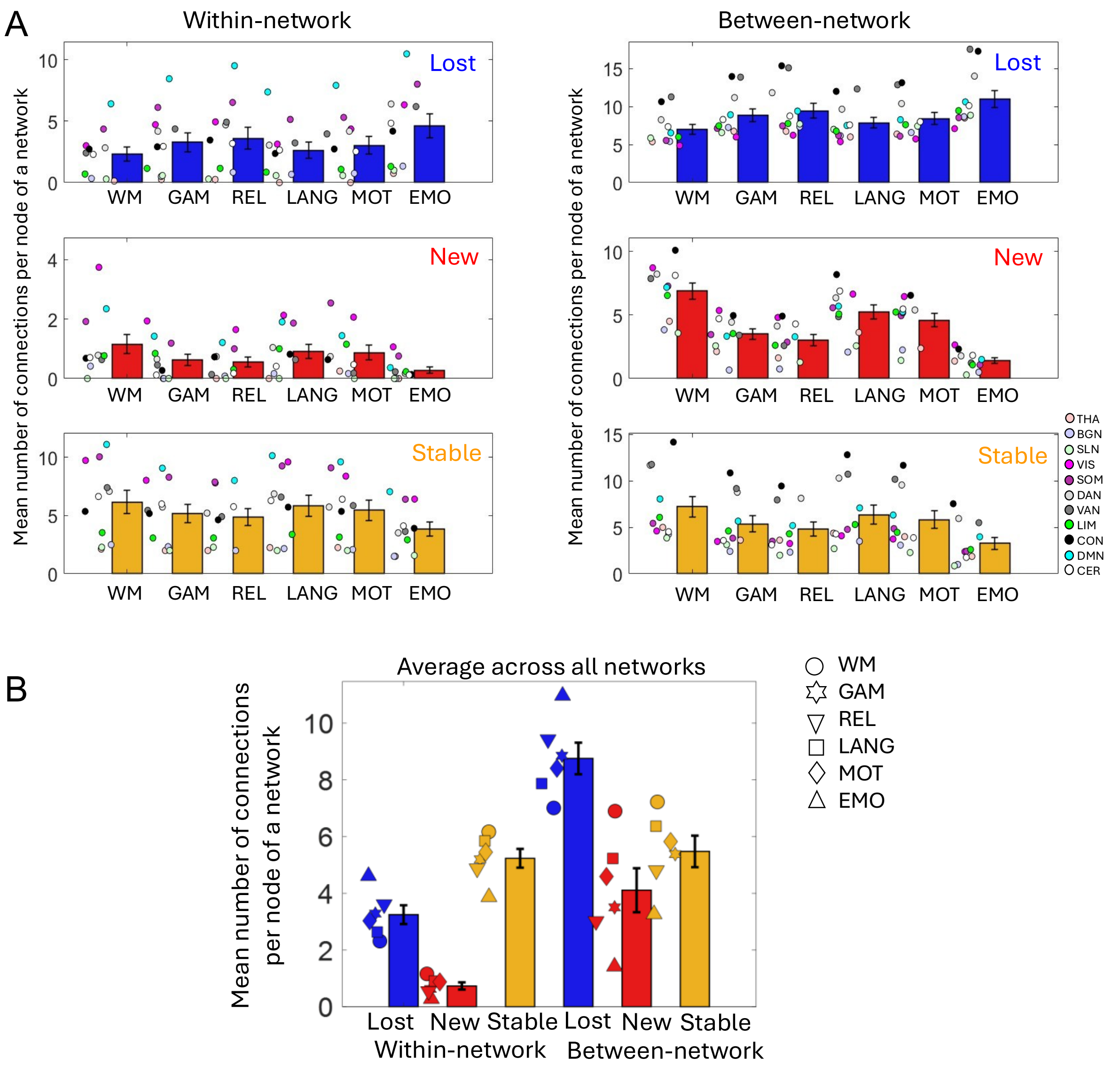


**Figure S1. Task-specific topological distribution of stable and reconfigured connections. (A)** Mean number of connections per node shown for each task—working memory (WM), gambling (GAM), relational processing (REL), language (LANG), motor (MOT), and emotion (EMO)—grouped by connection types (stable, newly formed, and lost) and network locations (within-network (WN) and between-network (BN)). These task-wise distributions underlie the averaged effects shown in **Fig. 2D**. Error bars denote standard error of the mean across 11 networks (thalamic (THA), basal ganglia (BGN), subcortical limbic (SLN), visual (VIS), somatomotor (SOM), dorsal attention (DAN), ventral attention (VAN), limbic (LIM), control (CON), default mode (DMN), and cerebellar (CER) networks). **(B)** Across all six connection categories (lost-WN, new-WN, stable-WN, lost-BN, new-BN, stable-BN), working memory and emotion task consistently occupied opposite ends of the distribution, with gambling, relational processing, language, and motor tasks showing intermediate profiles. While the overall pattern of stability vs. reconfiguration across network locations is conserved across tasks, individual tasks exhibit distinct modes, characterized by differential preservation, formation, and suppression of connections across network locations. Error bars indicate standard error of the mean across six tasks.

#### Statistical tables of repeated-measures ANOVA: Stability vs. Reconfiguration

**Table S1. Main and interaction effects of connection type and network location across tasks**

| **Task** | **Effect** | **Degrees of freedom** | **F-statistic** | **p-value** |
|---|---|---|---|---|
| WM | Connection type | 2, 20 | 18.80 | $2.5 \times 10^{-5}$ |
| | Location | 1, 10 | 23.50 | $6.7 \times 10^{-4}$ |
| | Connection type x Location | 2, 20 | 14.86 | $1.1 \times 10^{-4}$ |
| GAM | Connection type | 2, 20 | 30.75 | $7.9 \times 10^{-7}$ |
| | Location | 1, 10 | 18.23 | $1.6 \times 10^{-3}$ |
| | Connection type x Location | 2, 20 | 17.68 | $3.8 \times 10^{-5}$ |
| REL | Connection type | 2, 20 | 45.27 | $3.8 \times 10^{-8}$ |
| | Location | 1, 10 | 15.82 | $2.6 \times 10^{-3}$ |
| | Connection type x Location | 2, 20 | 20.91 | $1.3 \times 10^{-5}$ |
| LANG | Connection type | 2, 20 | 54.93 | $7.5 \times 10^{-9}$ |
| | Location | 1, 10 | 14.89 | $3.2 \times 10^{-3}$ |
| | Connection type x Location | 2, 20 | 21.78 | $9.5 \times 10^{-6}$ |
| MOT | Connection type | 2, 20 | 24.60 | $4.1 \times 10^{-6}$ |
| | Location | 1, 10 | 20.46 | $1.1 \times 10^{-3}$ |
| | Connection type x Location | 2, 20 | 16.25 | $6.4 \times 10^{-5}$ |
| EMO | Connection type | 2, 20 | 77.29 | $3.9 \times 10^{-9}$ |
| | Location | 1, 10 | 10.77 | $8.3 \times 10^{-3}$ |
| | Connection type x Location | 2, 20 | 30.81 | $7.8 \times 10^{-7}$ |

**Table S2. Post-hoc comparisons between network locations for connection types across tasks**

| **Task** | **Post-hoc comparisons** (Within − Between) | **Difference** | **Standard error** | **p-value** |
|---|---|---|---|---|
| WM | Lost | -4.70 | 0.80 | $1.6 \times 10^{-4}$ |
| | New | -5.74 | 0.58 | $1.7 \times 10^{-6}$ |
| | Stable | -1.05 | 1.31 | 0.44 |
| GAM | Lost | -5.38 | 0.98 | $2.6 \times 10^{-4}$ |
| | New | -3.72 | 0.44 | $8.0 \times 10^{-6}$ |
| | Stable | -0.37 | 1.11 | 0.75 |
| REL | Lost | -5.57 | 1.02 | $2.9 \times 10^{-4}$ |
| | New | -2.88 | 0.36 | $1.2 \times 10^{-5}$ |
| | Stable | -0.18 | 1.04 | 0.86 |
| LANG | Lost | -5.82 | 1.13 | $4.4 \times 10^{-4}$ |

| | | | | |
|---|---|---|---|---|
| | New | -2.47 | 0.34 | $2.9 \times 10^{-5}$ |
| | Stable | 0.07 | 0.95 | 0.95 |
| MOT | Lost | -5.23 | 0.91 | $1.8 \times 10^{-4}$ |
| | New | -4.32 | 0.48 | $3.8 \times 10^{-6}$ |
| | Stable | -0.52 | 1.17 | 0.67 |
| EMO | Lost | -6.36 | 1.30 | $6.2 \times 10^{-4}$ |
| | New | -1.14 | 0.19 | $1.1 \times 10^{-4}$ |
| | Stable | 0.60 | 0.78 | 0.46 |

### Task-specific topology of direct and indirect stable routes

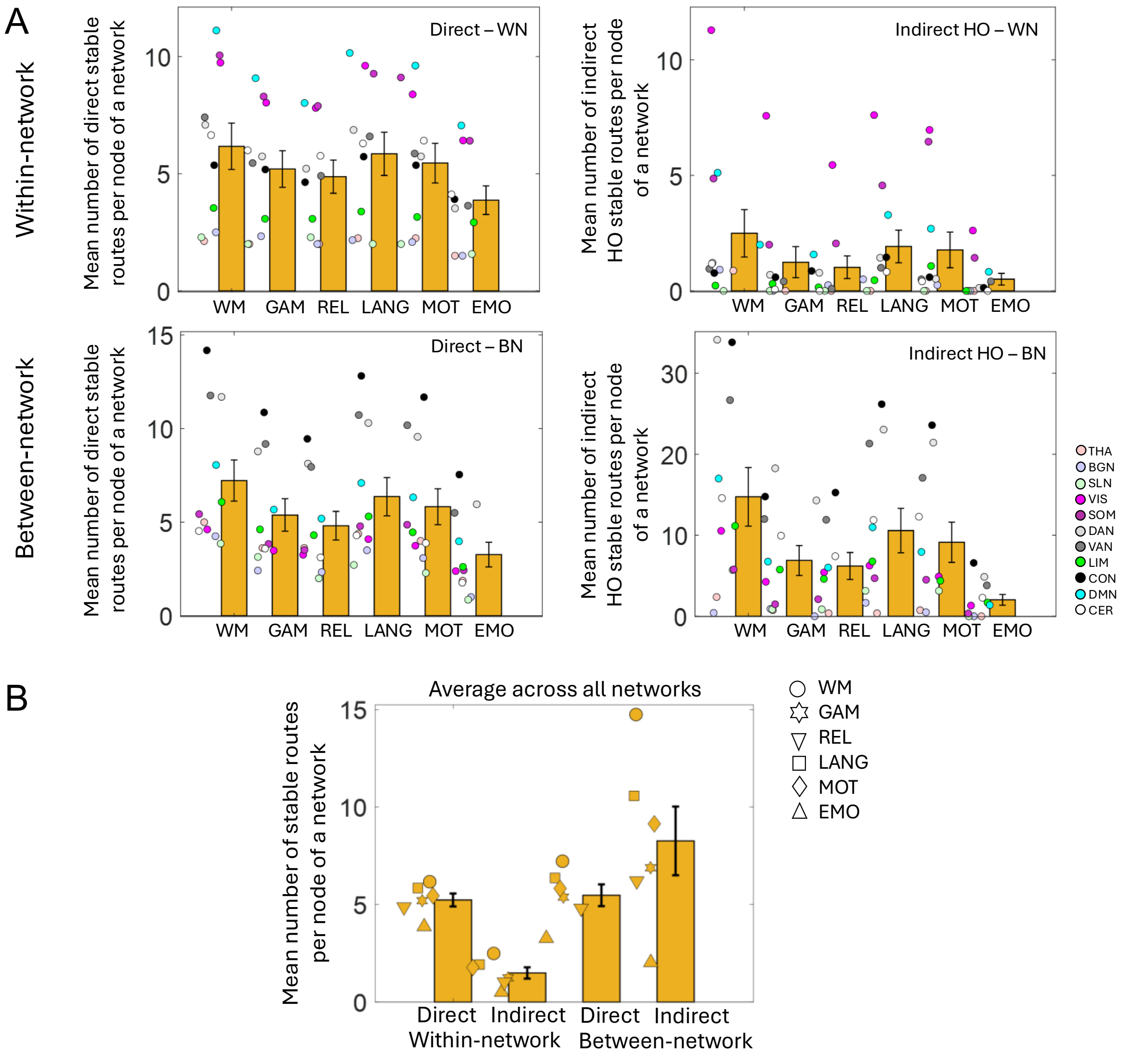


**Figure S2. Task-specific topological distribution of direct and indirect higher-order stable routes. (A)** Mean number of stable routes per node shown separately for each task, grouped by stable route types (direct and indirect higher-order (HO)) and network locations (WN, BN). These task-wise distributions underlie the averaged effects shown in **Fig. 3D**. Error bars indicate standard error of mean across 11 networks. **(B)** Across all four route categories (direct-WN, indirect HO-WN, direct-BN, indirect HO-BN), working memory consistently exhibited the highest mean number of stable routes, whereas the emotion task showed the lowest values, with gambling, relational processing, language, and motor tasks occupying intermediate ranges. Despite these task-specific differences in overall route abundance, the qualitative crossover pattern was preserved across tasks, with direct routes dominating within-network communication and indirect higher-order routes preferentially supporting between-network integration. Error bars indicate standard error of mean across six tasks.

#### Statistical tables of repeated measures ANOVA: Direct vs. Indirect Stable Routes

**Table S3. Main and interaction effects of route type and network location across tasks**

| Task | Effect | Degrees of freedom | F-statistic | p-value |
|---|---|---|---|---|
| WM | Route type | 1, 10 | 2.87 | 0.12 |
| | Location | 1, 10 | 7.42 | 0.02 |
| | Route type x Location | 1, 10 | 12.85 | $5.0 \times 10^{-3}$ |
| GAM | Route type | 1, 10 | 5.12 | 0.05 |
| | Location | 1, 10 | 4.23 | 0.07 |
| | Route type x Location | 1, 10 | 12.24 | $5.7 \times 10^{-3}$ |
| REL | Route type | 1, 10 | 7.29 | 0.02 |
| | Location | 1, 10 | 4.28 | 0.06 |
| | Route type x Location | 1, 10 | 16.36 | $2.3 \times 10^{-3}$ |
| LANG | Route type | 1, 10 | 0.04 | 0.85 |
| | Location | 1, 10 | 5.86 | 0.04 |
| | Route type x Location | 1, 10 | 13.78 | $4.0 \times 10^{-3}$ |
| MOT | Route type | 1, 10 | 0.08 | 0.79 |
| | Location | 1, 10 | 4.50 | 0.06 |
| | Route type x Location | 1, 10 | 11.03 | $7.7 \times 10^{-3}$ |
| EMO | Route type | 1, 10 | 64.12 | $1.2 \times 10^{-5}$ |
| | Location | 1, 10 | 0.38 | 0.54 |
| | Route type x Location | 1, 10 | 23.36 | $6.9 \times 10^{-4}$ |

**Table S4. Post-hoc comparisons between route types for network locations across tasks**

| Task | Post-hoc comparisons (Direct – Indirect) * | Difference | Standard error | p-value |
|---|---|---|---|---|
| WM | Within | 3.68 | 0.76 | $6.9 \times 10^{-4}$ |
| | Between | -7.52 | 2.62 | 0.02 |
| GAM | Within | 4.00 | 0.65 | $1.1 \times 10^{-4}$ |
| | Between | -1.50 | 1.79 | 0.23 |
| REL | Within | 3.85 | 0.51 | $2.1 \times 10^{-5}$ |

| | | | | |
|---|---|---|---|---|
| | Between | -1.4 | 1.00 | 0.19 |
| LANG | Within | 3.93 | 0.55 | $3.1 \times 10^{-5}$ |
| | Between | -4.21 | 1.80 | 0.04 |
| MOT | Within | 3.68 | 0.60 | $1.1 \times 10^{-4}$ |
| | Between | -3.32 | 1.66 | 0.07 |
| EMO | Within | 3.36 | 0.45 | $2.1 \times 10^{-5}$ |
| | Between | 1.24 | 0.25 | $5.3 \times 10^{-4}$ |

*Negative values indicate greater indirect than direct route counts

**Table S5. Post-hoc comparisons between network locations for route types across tasks**

| **Task** | **Post-hoc comparisons** (Within − Between) | **Difference** | **Standard error** | **p-value** |
|---|---|---|---|---|
| WM | Direct | -1.05 | 1.31 | 0.44 |
| | Indirect HO | -12.25 | 3.89 | 0.01 |
| GAM | Direct | -0.18 | 1.04 | 0.86 |
| | Indirect HO | -5.65 | 2.04 | 0.02 |
| REL | Direct | 0.07 | 0.95 | 0.95 |
| | Indirect HO | -5.19 | 1.73 | 0.01 |
| LANG | Direct | -0.52 | 1.17 | 0.68 |
| | Indirect HO | -8.65 | 2.86 | 0.02 |
| MOT | Direct | -0.37 | 1.11 | 0.74 |
| | Indirect HO | -7.37 | 2.77 | 0.02 |
| EMO | Direct | 0.60 | 0.78 | 0.46 |
| | Indirect HO | -1.53 | 0.74 | 0.07 |

#### Task-specific contribution of indirect higher-order routes to task-evoked information transfer

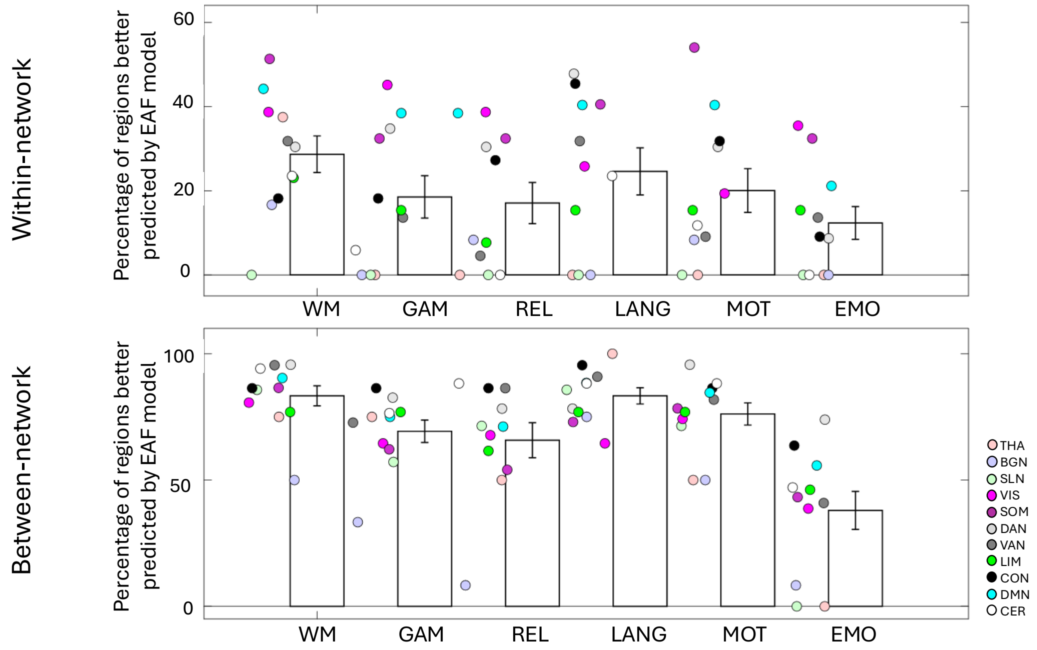


**Figure S3. Task-specific contribution of indirect higher-order routes to task-evoked information transfer.** Percentage of regions whose task-evoked activity was better predicted by the extended activity-flow model including indirect higher-order routes shown for each task, grouped by within-network (local) and between-network (global) levels. While the overall dominance of higher-order routes supporting task-evoked information transfer at the between-network level was conserved across tasks, distinct tasks exhibited systematic differences in overall percentage values. Working memory and language showed the highest percentage of regions benefiting from higher-order routes at both local and global scales, whereas emotion showed the lowest, with gambling, relational processing, and motor tasks occupying intermediate regimes (see **Supplementary Table S6** below). These task-wise distributions underlie the average effects shown in **Fig. 3E**. Error bars indicate standard error of mean across 11 networks.

#### Statistical table of paired comparisons performed between percentage values at network locations

**Table S6. Paired t-test performed for each task between percentage values at within- and between-network levels across 11 networks**

| Task | Within-network | Between-network | T-statistic (Within - Between) | p-value |
|---|---|---|---|---|

| | **Percentage of regions (mean(±SD) across networks)** | **Percentage of regions (mean(±SD) across networks)** | | |
|---|---|---|---|---|
| WM | 28.68 (±14.43) | 83.34 (±13.11) | -10.54 | $9.8 \times 10^{-7}$ |
| GAM | 18.54 (±16.70) | 69.30 (±14.73) | -9.06 | $3.9 \times 10^{-6}$ |
| REL | 17.08 (±16.25) | 65.77 (±22.97) | -6.08 | $1.2 \times 10^{-4}$ |
| LANG | 24.61 (±18.53) | 83.31 (±10.66) | -8.87 | $4.7 \times 10^{-6}$ |
| MOT | 20.06 (±17.23) | 76.15 (±14.61) | -11.91 | $3.1 \times 10^{-7}$ |
| EMO | 12.35 (±12.89) | 37.98 (±24.93) | -3.69 | $4.2 \times 10^{-3}$ |
